# Deep time structural evolution of retroviral and filoviral surface envelope proteins

**DOI:** 10.1101/2021.11.05.467493

**Authors:** Isidro Hötzel

## Abstract

The retroviral surface envelope protein subunit (SU) mediates receptor binding and triggers membrane fusion by the transmembrane subunit (TM). SU evolves rapidly under strong selective conditions, resulting in seemingly unrelated SU structures in highly divergent retroviruses. Structural modeling of the SU of several retroviruses and related endogenous retroviral elements with AlphaFold identifies a TM-proximal SU β-sandwich structure that has been conserved in the orthoretroviruses for at least 110 million years. The SU of orthoretroviruses diversified by differential expansion of the β -sandwich core to form domains involved in virus-host interactions. The β-sandwich domain is also conserved in the SU equivalent GP_1_ of Ebola virus although with a significantly different orientation in the trimeric envelope protein structure. The unified structural view of orthoretroviral SU and filoviral GP_1_ identifies an ancient, structurally conserved and evolvable domain underlying the structural diversity of orthoretroviral SU and filoviral GP_1_.

## Introduction

The retroviruses are an ancient group of viruses with wide genetic diversity. Of the three canonical retroviral genes, the *env* gene encoding the surface (SU) and transmembrane (TM) envelope protein subunits mediating receptor binding and membrane fusion during infection is the most diverse. TM is the more conserved of the two envelope protein subunits due to its role in membrane fusion during host cell infection. The SU subunit evolves more rapidly as it adapts to different hosts, receptors and during immune evasion. A long-standing question in retrovirus evolution has been the origin of the *env* gene of the *Retroviridae*^1–3^. In fact, it is not clear if the SU of retroviruses share any structural similarities or whether deep time evolution resulted in structurally distinct SU proteins without any remaining conserved structural elements.

The most biomedically important group of retroviruses are the orthoretroviruses. These include the alpharetroviruses, betaretroviruses, gammaretroviruses, deltaretroviruses, epsilonretroviruses and the lentiviruses which induce a variety of pathologies including oncogenesis and inflammatory and immunodeficiency syndromes in humans, mammals and avian species. Endogenous elements related to the orthoretroviruses are widespread in vertebrate genomes^4^. Most have large deletions and a heavy mutational load, especially in the *env* gene, that often leads to defective envelope proteins. However, orthoretroviral *env* genes integrated in vertebrate genomes can evolve under purifying selection and be maintained in an intact and sometimes functional form for millions of years. Among these are the syncytins, retroviral *env* genes coopted for key roles in placentation^4,5^.

Excluding the epsilonretroviruses, the envelope proteins of orthoretroviruses have been classified as beta-, gamma- and avian gamma-types^6,7^. The beta-type SU and TM envelope protein subunits betaretroviruses and lentiviruses are non-covalently associated and the TM subunit has 2 conserved cysteine residues^6,7^. The gamma-type envelope proteins of gammaretroviruses, deltaretroviruses have SU and TM subunits that are covalently associated, with a third conserved cysteine residue in TM mediating that association^6^. The avian gamma-type envelope protein is unique to the alpharetroviruses and is considered a variant of the gamma-type envelope^6^. The transmembrane subunits of orthoretroviruses are class I fusion proteins forming α-helical coiled-coils in the post-fusion state^8^. Other viral families with class I fusion proteins are the orthomyxoviruses, paramyxoviruses, coronaviruses, arenaviruses and filoviruses^8^. Sequence and structural similarities between the TM subunit of retroviruses and the GP_2_ transmembrane envelope protein subunit and filoviruses have been described^9,10^.

However, similarities in the receptor-binding envelope protein subunits of retroviruses and filoviruses are limited to the anchoring of the human immunodeficiency virus type 1 (HIV-1) gp120 and the Ebola virus GP_1_ surface envelope subunits to the transmembrane subunits through a pair of antiparallel β-strands^11^ but without any apparent sequence similarity.

The structural diversity of the SU of orthoretroviruses is apparent from the variety of domain organizations that are observed in different viral lineages. The SU of gammaretroviruses have a modular organization with an amino-terminal receptor-binding domain (RBD) followed by a proline-rich region (PRR) linking the RBD to a more conserved carboxy-terminal C-domain of unknown structure^12–16^. In contrast, HIV-1 gp120 does not have an independently-folding RBD analogous to the gammaretroviruses^17^. However, sequence and structural similarities are shared between of beta-type surface envelope proteins^18,19^. The regions of structural similarity correspond to a β-sandwich structure in the TM-proximal region of the inner domain of HIV-1 g120. Sections of this β-sandwich participate in intersubunit interactions with the gp41 transmembrane subunit^11,20^. The conserved β-sandwich structure is expanded differentially in the betaretroviruses and lentiviruses to form structurally distinct but topologically equivalent domains^21^. The apical regions of the betaretroviral SU models are topologically equivalent to the HIV-1 gp120 distal inner domain layer 2 that interacts with the coreceptor^22^. In contrast, the region topologically equivalent region to the HIV-1 gp120 β-sandwich expansion that forms the gp120 inner domain layer 3 and outer domain^22^ forms a short loop in the β-sandwich of the SU of betaretroviruses^21^. Thus, the beta-type SU proteins evolved by differential expansion of a conserved TM-proximal β-sandwich domain to form diverse structures mediating virus-host interactions.

It is not clear if the gamma-type SU, with its markedly different domain organization, also shares these or any other aspects of structural conservation with the betaretroviruses or lentiviruses. The recent release of AlphaFold 2, an artificial intelligence-based structural prediction tool that can yield structures of high quality comparable to experimentally determined structures^23,24^, provides an opportunity to address the structural conservation in the SU of retroviruses more broadly. Here is it shown that the structural diversity of orthoretroviral SU is based on differential expansion of an ancient and structurally conserved TM-proximal β-sandwich domain. In addition, this β-sandwich is also conserved in a structurally equivalent region of the SU equivalent GP_1_ of filoviruses, unifying a wide range of seemingly unrelated orthoretroviral and filoviral surface envelope proteins into a single protein family with well-defined structural properties.

## Results

### Modeling of orthoretroviral SU

A publicly available simplified version of AlphaFold 2^25^ was used to model the SU of 18 extant exogenous orthoretroviruses and endogenous retroviral elements (Fig. 1, Supplementary Figure S1). For simplicity, endogenous elements are classified here according to the orthoretroviral lineages from which they derive. Within the alpharetroviruses, the SU of the avian leukemia virus (ALV) and the lizard endogenous elements Mab-Env3 and Mab-Env4^26^ were modeled. In addition, the SU of a wide range of the more diverse gammaretroviruses and related endogenous elements were modeled. These include representatives of two major gammaretroviral subgroups differentiated by their transmembrane envelope protein sequences (Supplementary Fig. S1), named here type C and D. The type C gammaretroviruses include the murine (MLV) and feline (FELV) leukemia viruses and porcine (PERV)^27^ and python (PyERV)^28^ endogenous retroviruses. The type D gammaretroviruses include the avian reticuloendotheliosis virus (REV) and the endogenous baboon (BaEV) and feline RD114 endogenous retroviruses. The Mason-Pfizer monkey virus (MPMV) is a betaretroviruses that acquired a gamma-type *env* gene by recombination from a virus related REV^29^ and is classified here as a type D gammaretrovirus based on its envelope protein sequence (Supplementary Fig. S1). In addition, models were also obtained for the SU of human syncytin-1 and 2^30,31^, which cluster by TM sequence with the type D gammaretroviruses, the unclassified gammaretroviral mammalian syncytin-Mar1^32^, syncytin-Car1^33^ and syncytin-Rum1^34^ and the SU encoded by endogenous gammaretroviral elements from spiny-rayed fishes^35^. The latter include the SU encoded by the mudskipper *percomORF* retroviral *env* endogenous element and a related retroviral endogenous element from the Japanese Eel, named here Env-Psc and Env-Aja.Modeling of the SU of additional endogenous gammaretroviral elements, deltaretroviruses and epsilonretroviruses was unsuccessful, yielding mostly unfolded models that were not further analyzed.

**Figure 1.**
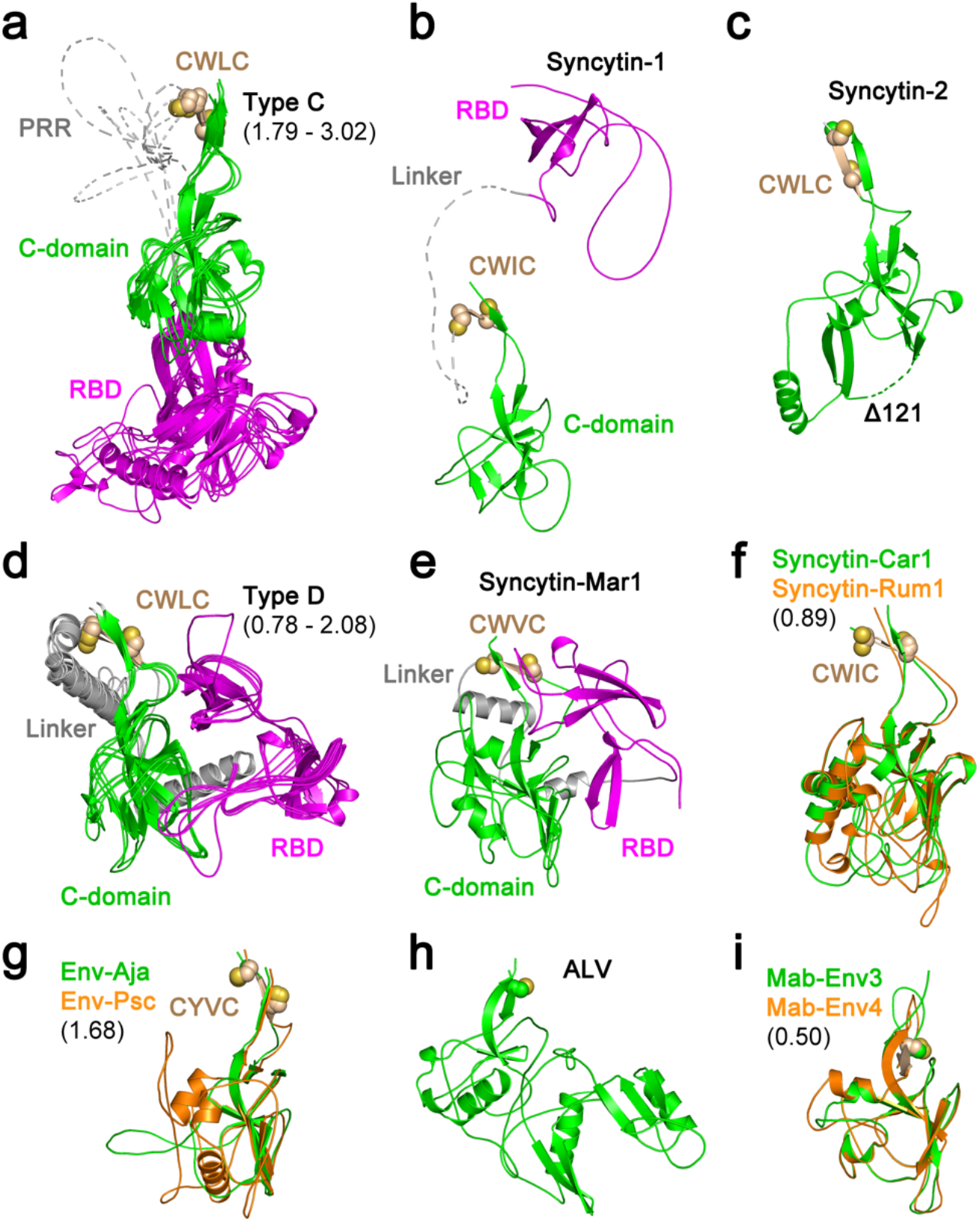
Orthoretroviral SU structural models. Models of the SU of (a) type C gammaretroviruses, (b) syncytin-1, (c) syncytin-2, (d) type D gammaretroviruses, (e) syncytin-Mar1, (f) syncytin-Car1 and Rum1, (g) Env-Aja and Env-Psc, (h) ALV and (i) Mab-Env3 and Mab-Env4. Panels a, d, f, g and i show superpositions of structurally similar models with minimum and maximum root mean square deviation of aligned models in Angstroms shown in parentheses. The models in panels a, b, d and e show the RBD in magenta and the linker regions joining the RBD to the C-domain in gray. The models in panels f, g and i are shown in colors as indicated in each panel. The disordered PRR and linker regions are shown as dotted lines in panels a and b. The conserved cysteine residues in the gammaretroviral CWLC consensus motif in panels a to g or preceding the first β-strand in the alpharetrovirus SU models in panels h and i are shown as spheres. The deletion in the syncytin-2 SU model (122) is shown as a dotted line.

A high diversity of modeled SU structures was observed, with SU models clustering according to TM sequence similarities in most cases (Fig. 1, Supplementary Fig. S1). SU models had predicted Local Distance Difference Test (pLDDT) scores of at least 70 and often above 90, which indicate high-confidence main chain modeling^23^ (Supplementary Fig. S2, S3). The main exceptions were relatively short sections in the SU termini, which had lower pLDDT scores and were mostly unfolded (Supplementary Fig. S2). These regions were excluded from the analyses. Some models had relatively long internal regions with low pLDDT scores (Supplementary Fig. S2). Many of the low-scoring internal regions correspond to long linker regions between domains (Fig. 1a, b, d, e, Supplementary Fig. S3a-e). For syncytin-2 SU, a high-scoring model similar to SU models of other gammaretroviruses was obtained only after deleting 122 residues from its mid-section (Fig. 1c, Supplementary Table S1). Thus, the syncytin-2 SU structure modeled here is significantly different from the syncytin model in the AlphaFold Protein Structure Database^23^ without the deletion (UniProt accession P60508), which has generally lower-quality pLDDT scores in the SU region. The quality of the SU models obtained allows the comparison of domain organization and overall secondary and tertiary structural elements among a wide range of orthoretroviruses.

Two major SU groups with different amino-terminal RBDs were identified within the gammaretroviruses (Fig. 1a, b, d, e). One includes the SU of type C gammaretroviruses with an RBD similar to the previously described MLV and FELV RBD crystal structures^15,16^. The other group includes the type D gammaretroviruses as well as the SU of human syncytin-1 and rodent syncytin-Mar1. In neither group the position of the RBD relative to other domains of the models may be reliable due to the long and apparently flexible linkers between domains. In fact, The type D gammaretroviruses have an amino-terminal RBD that consists of two topologically similar β-sheets (Fig. 2a, b). Syncytin-1 and syncytin-Mar1 SU models have RBDs structurally similar to the carboxy-terminal subdomain of the REV RBD model despite little to no sequence similarity (Fig. 2b, c, d). The similarities between the RBD structures of syncytin-1 and the type D gammaretroviruses is consistent with syncytin-1 sharing the same receptor as RD114 and MPMV^36^. No significant structural similarities were observed between the RBDs of type C and D gammaretroviruses.

**Figure 2.**
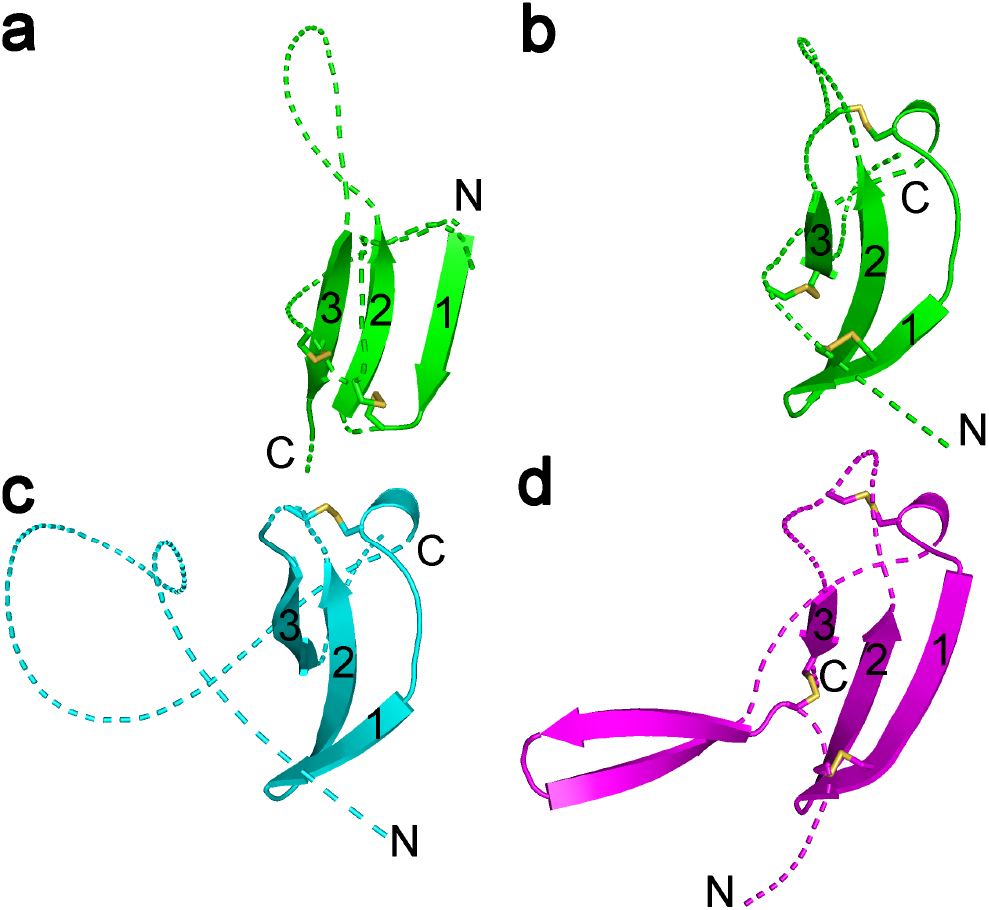
RBD of type D gammaretroviruses, syncytin-1 and syncytin-Mar1. (a) First, amino-terminal β-sheet of the type D REV RBD. (b) second, carboxy-terminal REV RBD β-sheet. (c) Syncytin-1 RBD. (d) Syncytin-Mar1 RBD. The β-sheets are shown in the same orientation in all panels with the three sequential β-strands indicated by numbers. Loops linking β-strands and flanking the β-sheet region are shown as dotted lines for clarity. Disulfides are shown as sticks. The amino and carboxy termini of each section are indicated.

An independently-folding carboxy-terminal domain in the SU of type C and D gammaretroviruses, syncytin-1 and synctin-Mar1 corresponds to the entire C-domain. This domain starts with a β-strand including the conserved gammaretroviral CWLC consensus motif that mediates intersubunit covalent interactions^37,38^ (Fig. 1a-g). Surprisingly, the SU models of a subset of unclassified gammaretroviral elements are comprised of a C-domain structure without an independent RBD, including the SU of the type D-like gammaretroviral syncytin-2 (Fig. 1c, f, g). A CWLC-like consensus motif β-strand in these models is located near the SU amino terminus. Several of these gammaretroviral syncytins are fusogenic^30,32–34^, indicating that receptor binding is mediated by the C-domain equivalents of these envelope proteins. The models for the SU of alpharetroviruses do not have an independent amino-terminal RBD (Fig. 1h, i). The alpharetroviruses do not have a CWLC-like motif but the first β-strand is preceded by a conserved cysteine residue (Fig. 1h, i). In the Mab-Env3 SU model this cysteine forms a disulfide with an amino-terminal cysteine residue in an unstructured region outside the main domain, while in the ALV and Mab-Env4 SU models these cysteines are unpaired.

### Identification of a conserved orthoretroviral SU structure

Despite the structural diversity in the SU models of different retroviral groups, a highly conserved domain shared by all orthoretroviral lineages was identified (Fig. 3, 4). This domain comprises the C-domain of type C and D gammaretroviruses and the entire SU of unclassified gammaretroviruses without an RBD and alpharetroviruses. The conserved domain corresponds to the gp41-proximal HIV-1 gp120 β-sandwich^17,22^ (Fig. 4a), also conserved in the SU of other lentiviruses and betaretroviruses^21^. A structural alignment of the models excluding the amino-terminal RBD of gammaretroviruses defined 12 consensus β-strands, 10 of which have homologues in the HIV-1 gp120 proximal inner domain structure (Fig. 3). This domain thus represents a highly conserved TM-proximal domain (PD) of orthoretroviral SU. To harmonize the nomenclature of regions in the different models and structures, the consensus β-strands of the PD are numbered 1 to 12 and connecting regions denoted sequentially from A to K (Fig. 3, 4, Supplementary Fig. S4). The region homologous to the region that in HIV-1 gp120 faces the trimer axis is named “layer 1” according to its designation in HIV-1 gp120^22^. This region is structurally diverse and has lower pLDDT scores in several models (Supplementary Fig. S3). All cysteines in the PD except those in the terminal β-strand 1 of the gammaretroviruses are involved in disulfide bonds in the models but with some notable differences in the arrangement of disulfide bonds by structurally equivalent cysteines in different viral lineages (Fig. 3, 4). Although distinct, all disulfides are compatible with the conserved β-sandwich fold. For reasons that are unclear, four cysteines in the type C gammaretroviral PD models form disulfides that are distinct from the experimentally determined disulfides in the SU of MLV and the related mink cell focus-forming virus^39,40^. The experimentally determined disulfides would not be compatible with the conserved PD β-sandwich fold. However, in alpharetroviral SU, receptor binding is sufficient to disrupt a disulfide between the β-strands 8 and 11 homologues^41^, suggesting that disulfide bonds may be relatively labile in that region. The conserved gammaretroviral CWLC consensus motif in β-strand 1 is expected to be located nearby the TM cysteine that covalently links with SU by analogy with the HIV-1 pre-fusion trimeric envelope crystal structure^11^.

**Figure 3.**
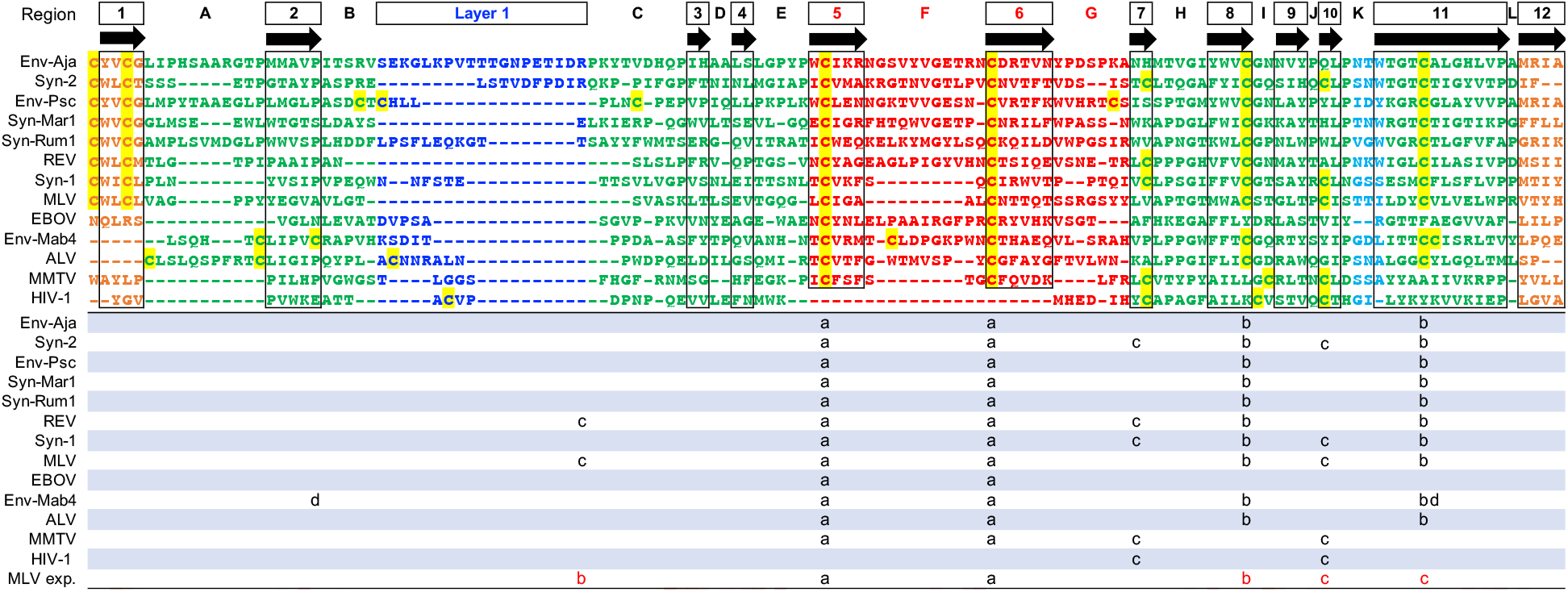
Conserved orthoretroviral and filoviral PD. Orthoretroviral and filoviral PD sequences structurally aligned to the PD of Env-Aja. Insertions relative to Env-Aja SU are not shown. Only one representative of each orthoretroviral and filoviral lineage is shown. The consensus β-strands are indicated in boxes and arrows. Cysteine residues are highlighted in yellow. Disulfide bonds in models and structures are shown below the structural alignment, with the last line indicating the experimentally determined disulfide bonds in MLV SU. The proximal domain is highlighted in green except for the apical and layer 1 regions, highlighted in red and blue. The terminal β-strands 1 and 12 are highlighted in orange.

**Figure 4.**
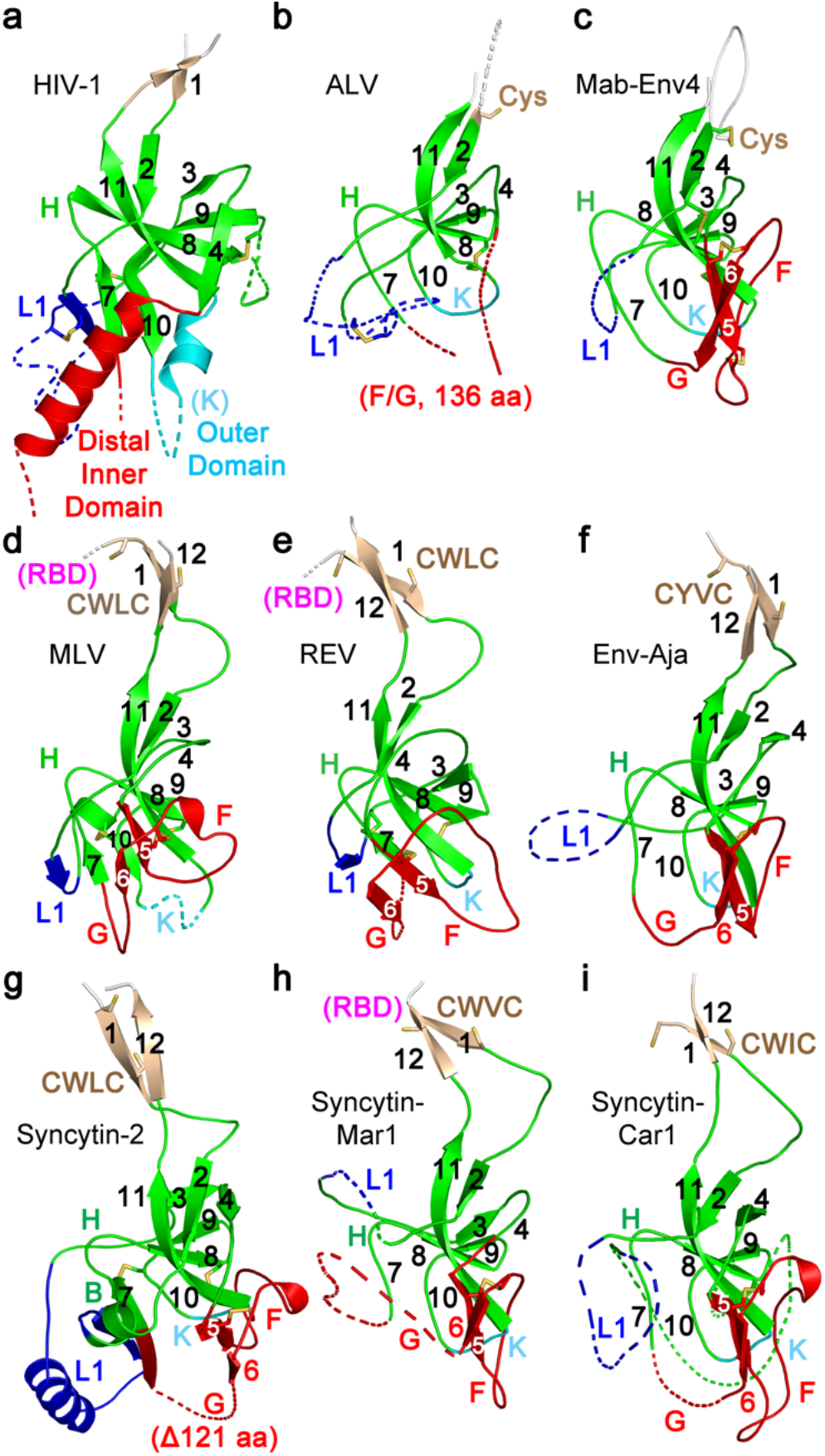
Conserved orthoretroviral SU proximal domain β-sandwich. (a) Structure of HIV-1 gp120 PD region (PDB 3JWD). PD regions of the (b) ALV, (c) Mab-Env4, (d) MLV, (e) REV, (f) Env-Aja, (g) syncytin-2, (h) syncytin-Mar1 and (i) syncytin-Car1 SU models. Parts of layer 1 (L1) and regions F, G and the HIV-1 gp120 distal inner domain and outer domain are shown as dotted lines for clarity. The PD, apical domain and layer 1 regions are shown in green, red and blue. Selected loop and β-strand regions are labeled in each panel. Cysteine residues are shown as sticks. The location of the conserved CWLC consensus motifs of gammaretroviral SU in β-strand 1 is indicated in panels d to i. The connections to the RBD in the models in panels d, e and h are shown in magenta. All structures and models shown in the same orientation as in Fig. 1. Dotted lines indicate regions omitted for clarity.

Structural conservation of the β-sandwich is observed despite absence of any significant sequence similarity between orthoretroviral groups within the region (Fig. 3). The structural conservation includes the complex topological arrangement of β-strands and their connections within the β-sandwich (Fig. 3, 4, Supplementary Fig. S4). In all models the chain between β-strands 2 and 7 circles around the conserved domain in the same clockwise direction as observed from the virion surface (Fig. 4, Supplementary Fig. S4). The β-sheet formed by antiparallel strands 8, 9 and 11 have β-strand 8 centrally located between the other two β-strands. The two β-strands that are observed in the PD of all SU models but not in HIV-1 gp120 are parallel β-strands 5 and 6, which extend the β-sheet formed by antiparallel β-strands 9, 8 and 11 (Fig. 3, 4, Supplementary Fig. S4). These two β-strands are joined in a right-handed configuration by disulfide-constrained loop F which wraps around the beginning of β-strand 11 in most models except in the SU of type C gammaretroviruses with a shorter loop F. Together with loop G, these form an apical region in the conserved PD. Analysis of the previously described betaretroviral SU models^21^ identified β-strand 5 and 6 homologues, including the conserved disulfide, and region F and G homologues. Thus, the SU of lentiviruses is unique in that the region corresponding to the apical region of orthoretroviral SU forms an α-helix that projects towards the distal end of the inner domain^17,21^ rather than a conserved loop.

Structural alignments with significant DALI similarity Z-scores^42^ were obtained between most SU models (Fig. 5). The exceptions are the HIV-1 gp120 crystal structure and the betaretroviral mouse mammary tumor virus (MMTV) SU model, which had the generally lowest similarity scores and failed to align with some SU models. The highest average structural similarity scores were observed for the SU models of human syncytin-2 and the unclassified gammaretroviral Env-Aja endogenous element. These SU models have PD structures that are the closest to the consensus of all models. The SU models with the highest DALI Z-scores when aligned with the syncytin-2 SU model were derived from Env-Aja followed by the type D gammaretroviruses, gammaretroviral syncytins, the Env-Psc *percommORF* endogenous element, ALV and the related alpharetroviral lizard syncytins Mab-Env3 and Mab-Env4, type C gammaretroviruses and betaretroviruses and finally the crystal structure of HIV-1 gp120 (Fig. 5). The conservation of very specific structural features indicate a common origin for the PD of different orthoretroviral lineages.

**Figure 5.**
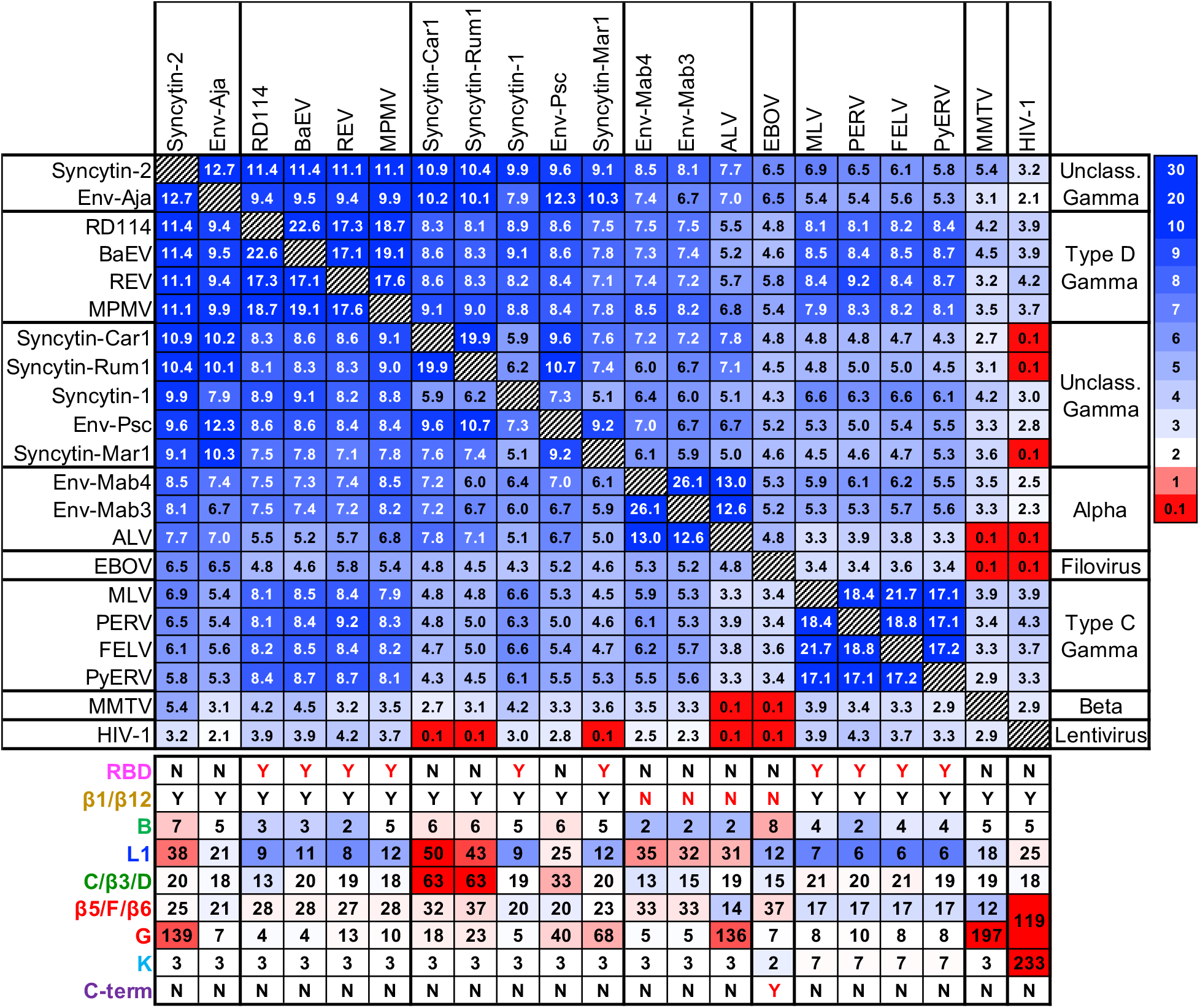
Orthoretroviral SU and filoviral GP_1_ DALI structural similarity Z-score matrix. The viral lineages and the scale for DALI Z-scores are shown on the right. The length in amino acids of selected PD regions of different viruses and endogenous elements are shown below the matrix. Color coding indicates lengths below (blue tones), above (red tones) or equal to the median for each region.

### Differential expansion of the PD in the orthoretroviruses

The SU of orthoretroviruses is formed in most cases by expansions and extensions of the PD as it has been observed for the betaretroviruses and lentiviruses but with notable differences among and within orthoretroviral lineages. The PD of the type C and D gammaretroviruses does not have long internal expansions analogous to the betaretroviruses and lentiviruses (Fig. 4d, e). Instead, the major expansion of the PD of type C and D gammaretroviruses, syncytin-1 and syncytin-Mar1 is the amino-terminal linker and RBD (Fig. 1a, b, d, e, 4d, e ,h). As mentioned earlier, the SU of unclassified gammaretroviral endogenous elements Env-Aja, syncytin-2, syncytin-Car1, syncytin-Rum1 and Env-Psc do not have a domain comparable to the type C and D gammaretroviral RBDs (Fig. 1c, f, g). Instead, syncytin-2 SU has a long expansion of loop G that is topologically equivalent to the betaretroviral SU apical region (Fig. 4g). Syncytin-Car1, syncytin-Rum1 and Env-Psc SU are more compact, with smaller expansions of different subsets of layer 1, the section from regions C to D and the apical loop G (Fig. 1f, g, 4h, i). The Env-Aja SU is comprised of a PD with no significant extensions or expansions (Fig. 1g, 4f). The minimalistic SU domain organization of Env-Aja combined with its near-consensus PD structure indicates that it is closely related structurally to a basal orthoretroviral SU.

The avian gamma-type alpharetroviral ALV SU model has a domain organization that is analogous to the SU of betaretroviruses and syncytin-2 (Fig. 4b). Similarities include a relatively long expansion of apical region F/G but not region K. The alpharetroviral-like syncytins Mab-Env3 and Mab-Env4 have PD structures that most closely resemble the PD of ALV SU but without any extensions of region G or any other region relative to ALV SU (Fig. 1i, 4c, Fig. 5). These two endogenous elements, like the Env-Aja SU, also have a minimalist SU comprised of a PD without significant extensions except for a slightly longer loop F and layer 1 relative to Env-Aja (Fig. 5).

### Conservation of the PD in filoviral GP_1_

The structural similarities among the SU of distantly related orthoretroviruses can be observed despite any obvious similarities in the underlying sequences. Given the sequence and structural similarities between the TM subunit of retroviruses and transmembrane envelope protein GP_2_ of filoviruses, it is possible, although it has not been reported, that these structural similarities extend to filoviral GP_1_. Visual inspection of Ebola virus (EBOV) trimeric envelope protein crystal structures^43,44^ revealed that the virion-proximal region of the GP_1_ subunit including the base and head subdomains has a fold similar to the conserved orthoretroviral PD β-sandwich (Fig. 6a, Supplementary Fig. S5). The topology of the chain in the filoviral PD homologue is the same as for the orthoretroviral PD. Structural similarities include the clockwise turn of the chain between the β-strands 2 and 7 homologues around the domain and the relative positions of the antiparallel β-strands 8, 9 and 11 homologues. In addition, the apical region formed by the β-strands 5 and 6 and loop F homologues of EBOV GP_1_ is structurally similar to orthoretroviral SU, including the conserved disulfide (Fig. 3, 6a). Differences include a slightly expanded region B, a slightly longer loop F relative to type C and D gammaretroviral SU, Mab-Env3 and Mab-Env4 and continuous β-strands 1/2 and 11/12 in the proximal region in EBOV GP_1_. Structural alignments between EBOV GP_1_ and the SU of orthoretroviruses were significant in all cases except for the SU of MMTV and HIV-1 gp120, with the highest similarity scores for the alignments with the consensus Env-Aja and syncytin-2 model structures (Fig. 5). The glycan cap and mucin-like domains that are cleaved prior to binding of GP_1_ to the receptor in the endosome_43,45_ are thus carboxy-terminal extensions of the conserved β-sandwich structure (Supplementary Fig. S4).

**Figure 6.**
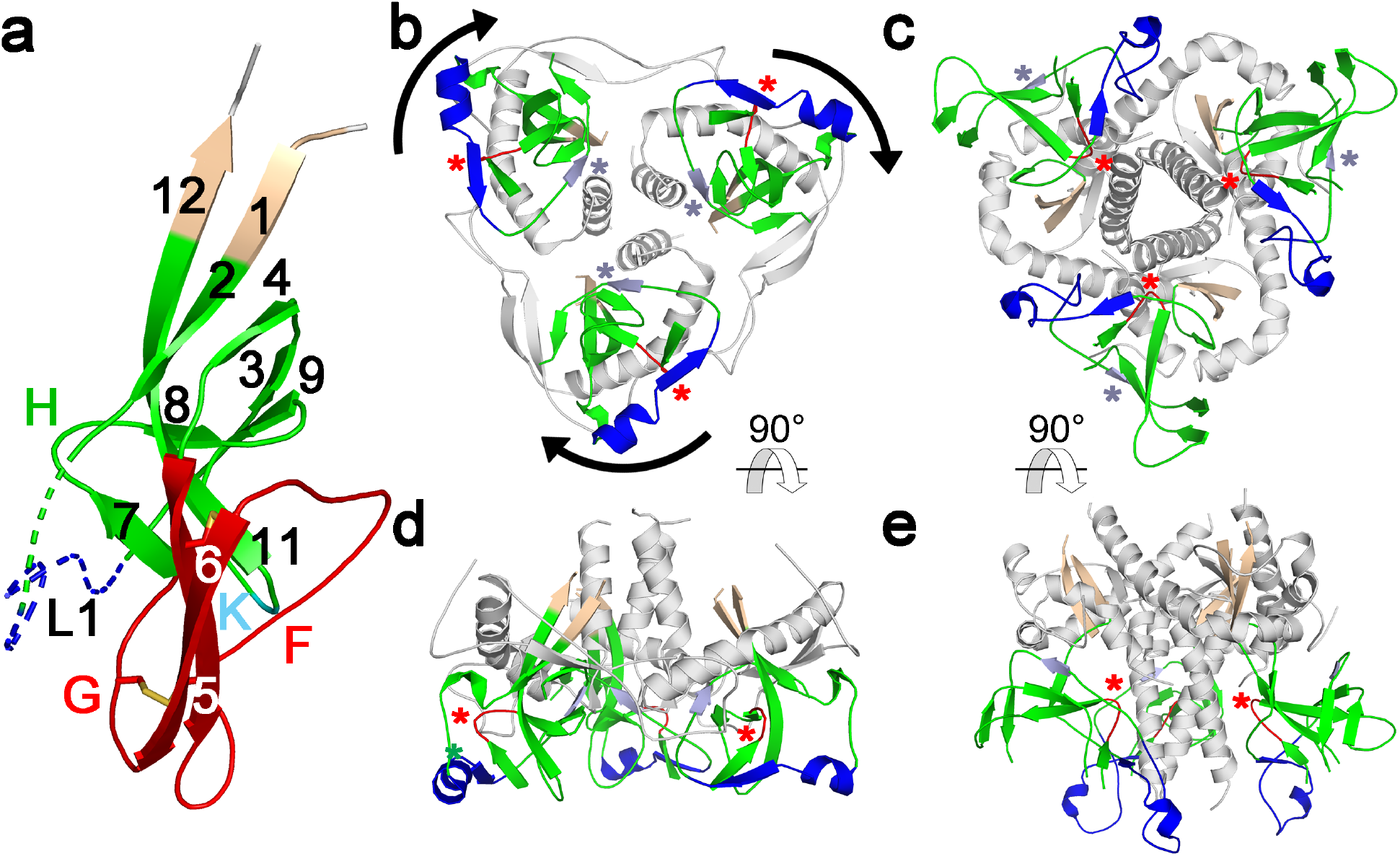
Conservation and orientation of the PD in EBOV GP_1_. (a) EBOV GP_1_ base and head subdomains comprising the PD domain (PDB 3CSY), shown in a similar orientation as the PD of orthoretroviruses in Fig. 4. The β-strand and selected loop regions are labeled and colored as in Fig. 2 and 3. Disulfides in the apical region are shown as sticks. (b) EBOV (PDB 5HJ3) and (c) HIV-1 (PDB 4TVP) trimeric envelope protein crystal structures shown from the top, distal side. The apical domain in the GP_1_ protomers in panel b and the distal region of the inner domains and the outer domains of gp120 in panel c are omitted for clarity. The β-sandwich, terminal β-strands and layer 1 homologues are colored green, wheat and blue. Region H and β-strand 3 homologues are colored red and light blue and highlighted with red and light blue asterisks. The arrows in panel b indicate the clockwise direction of the major 120° rotation of GP_1_relative to gp120 protomers in the trimeric structures. Panels (d) and (e) show the same structures as panels b and c, rotated 90°, with the virion-proximal side facing up.

The most remarkable difference between the HIV-1 gp120 and EBOV GP_1_ structures is the orientation of the β-sandwich PD in the envelope trimers. The EBOV GP_1_PD is rotated clockwise by approximately 120° around the axis perpendicular to the viral membrane compared to the HIV-1 PD as seen from the top of the trimer (Fig. 6b, c). This results in different sets of PD regions occluded in the envelope protein trimers of HIV-1 gp120 and EBOV GP_1_. For instance, whereas region H and β-strand 3 in the gp120 protomers of the HIV-1 trimeric envelope protein crystal structure are facing the trimer axis and the surface of the trimer respectively, the opposite is true for the homologous regions of GP_1_ protomers in the EBOV trimeric envelope protein crystal structure (Fig. 6b, c, d, e). None of the intersubunit contacts mediated by residues in the PD and terminal β-strands are structurally equivalent in these two viruses (Supplementary Fig. S6).

## Discussion

The structural models presented here provide a unified view of the structure of the SU of most biomedically important orthoretroviruses and related endogenous elements that play critical roles in placentation. This is likely to be extend to the deltaretroviruses and all lentiviruses for which SU structures or reliable models have not yet been obtained. Furthermore, the same basic structural type is also observed in the GP_1_ protein of filoviruses, thus unifying the envelope proteins of these different viral families, including the receptor-binding subunits, into a single structural protein family. AlphaFold consistently yields models with similar features despite the absence of sequence similarity between different viral lineages in the regions modeled. Thus, collectively, the number of models converging to similar structures indicate robust and reliable models in addition to the relatively high pLDDT reliability scores, especially in the PD region. Some regions are poorly modeled, as for example the apical regions of syncytin-2 and ALV SU and perhaps the type D gammaretroviral RBDs. The overall domain organization of the SU of different orthoretroviruses can nonetheless be gleaned from the models to identify the main shared structural features and the structural evolution of these proteins.

The major insight that emerges from the unified structural view is that at the center of the highly diverse and rapidly evolving receptor-interacting surface envelope protein subunit sits a structurally conserved but evolvable domain, the PD β-sandwich. Sequence variation is tolerated within the PD β-sandwich while retaining its basic structure. At the same time, the PD allows large expansions that emanate from several loops within the domain or its termini, resulting in a diversity of SU and GP_1_ structures with important functions in receptor interaction and immune evasion in different viral lineages.

A second insight emerging from this unified structural view is the age of the conserved SU β-sandwich structure of orthoretroviruses and filoviruses. The oldest known members of the orthoretroviral family with the conserved PD structure are the spiny-rayed fish *percomORF* endogenous elements. The *percomORF* element genomic integration event was dated to at least 110 million years ago^35^, setting a lower-bound estimate for the age of the conserved orthoretroviral PD structure. The retroviral element encoding Env-Aja has not been dated but its *env* gene seems to be basal to the *percomORF* elements by TM protein sequence analysis^35^ and may therefore derive from an even older orthoretroviral lineage. The ultimate origin of the orthoretroviral *env* gene with its conserved SU is unresolved. High reliability score models of the SU of spumaretroviruses, the ancient sister group of orthoretroviruses within the retroviruses^46,47^, have no structural similarity with the orthoretroviral SU models (Supplementary Fig. S7), indicating a different origin for the *env* gene of these two major retroviral lineages. It is possible that orthoretroviruses acquired an *env* gene from another viral lineage early in their evolution, similar to *env* gene acquisition by LTR retrotransposable elements of invertebrates^1–3^. One possible source is a filovirus-like envelope protein gene. However, due to the antiquity of orthoretroviral SU, it is possible that a filovirus-like lineage acquired its envelope protein gene from an orthoretrovirus with an SU structurally similar to that of Env-Aja, Mab-Env3, Mab-Env4 or an unknown or extinct orthoretroviral lineage with a C-terminal extension in the PD. A similar transfer based on sequence similarities between orthoretroviral TM and GP_2_ was previously suggested^10^. The definition of the basic shared structural elements of this conserved domain as well as the patterns of sequence variation in the structural models described here and in endogenous elements in genomic data from multiple vertebrate species may allow identification of more distantly related members of this protein family.

The unified structural view also provides information about the structural evolution of SU in different orthoretroviral lineages. The structural models indicate that divergence of alpha and gammaretroviruses occurred before or shortly after the start of structural diversification by PD expansion. Both the alpha and gammaretroviruses include endogenous elements encoding SU proteins consisting of minimal or almost minimal PD structures without an amino-terminal RBD or other major expansions. The most parsimonious interpretation of the data is that the minimalistic SU of Env-Aja, Mab-Env3 and Mab-Env4 are closely related structurally to a basal orthoretroviral SU and that most or all of structural diversification by terminal and internal expansions of the PD occurred independently and in parallel after divergence of these lineages. For the beta-type envelope proteins it is not clear if SU structural diversification occurred before or after divergence of the betaretroviruses from the lentiviruses as no beta-type minimal SU structures have been identified. Lastly, the SU of lentiviruses is the most divergent structurally within the orthoretroviruses and filoviruses, with one or more outer domains derived from region K and a structurally unique distal inner domain in place of the PD apical domain^17,21^.

An unexpected source of structural variation in this protein family is the orientation of the PD in oligomeric envelope protein structures. This results in different faces of the PD interacting with the transmembrane subunits in HIV-1 and EBOV trimeric envelope proteins. The only structurally equivalent interactions of the PD of HIV-1 and EBOV with TM and GP_2_ are the terminal regions of the domain outside the β-sandwich^11^. Not only is the PD oriented differently in these viruses but TM and GP_2_ also fold differently and make contacts with the PD that are not structurally analogous in the respective pre-fusion structures. Therefore, the intersubunit interface seems to be evolutionarily malleable, with the terminal strand attachment of the PD to the transmembrane subunit being the only interaction that remained structurally conserved but with some degree of structural variation. The difference in the orientation of the EBOV and HIV-1 PD in the trimeric structures is not analogous to the rotation that HIV-1 gp120 undergoes within the trimer upon receptor binding^48,49^. Glycosylation site patterns suggest that the PD of extant gammaretroviruses may adopt an orientation different from the PD of HIV-1 in trimeric envelope proteins. The HIV-1 PD surface that interacts with the trimer axis is devoid of glycosylation sites^17,20^. In contrast, the homologous PD surface in extant gammaretroviruses often has glycosylation sites, including in region H which in the HIV-1 envelope protein trimeric structures is closely associated with TM (Supplementary Fig. S8), suggesting that it does not interact with TM. The gammaretroviral PD surface equivalent to the EBOV PD surface interacting with GP_2_ is mostly devoid of glycosylation suggesting that the gammaretroviral PD orientation in trimeric structures is similar to the EBOV PD or intermediate between filoviruses and lentiviruses.

The differences in PD orientation and intersubunit interactions mediated by the β-sandwich structure of HIV-1 gp120 and EBOV GP_1_ suggest that the conservation of this structure is not primarily due to the preservation of critical intersubunit interactions that maintain the envelope protein trimer in its metastable pre-fusion state prior to receptor binding. Other functional activities of the PD probably account for its deep time structural conservation. Spacing requirements between the cell surface and the transmembrane subunits for optimal membrane fusion may be met by PD expansions rather than the β-sandwich structure itself. It is possible that the events after receptor binding that release TM and GP_2_ in an optimal manner for membrane fusion are mediated by conformational changes within the PD that depend on its conserved tertiary structure. Alternatively, receptor binding may lead to a rigid body movement of the PD to trigger membrane fusion that depends on the geometry of receptor binding relative to the terminal β-strand intersubunit interactions.

The conserved apical region structure may be a critical element in the mechanism of receptor-induced membrane fusion. Interaction of the PD apical region with a receptor appears to be a shared functional event to trigger membrane fusion in this protein family. This has been shown for the filoviruses, alpharetroviruses and even for the type C gammaretroviruses, which depend on sequences that include the PD apical region to induce fusion after receptor binding by the RBD^12–15,45,50,51^. HIV-1 gp120 interacts with its coreceptor through a structurally distinct but topologically equivalent region, the distal region of the inner domain, that connects with the rest of the PD in a manner similar to the apical region of other viral lineages^17^. The minimal orthoretroviral SU variants identified here may serve as simple and structurally tractable systems to elucidate the basic functions of the conserved PD structure without the layers of structural complexity used by extant orthoretroviruses and filoviruses for receptor binding and immune evasion.

## Methods

### Structural modeling

The mature orthoretroviral SU sequences (Supplementary Table S1) were used as input for structural modeling with AlphaFold 2^23^. The accession numbers for SU sequences are shown in Supplementary Table S1. A simplified version^25^ of Alphafold 2 run on the ColabFold server with graphical processing unit support accessible at https://colab.research.google.com/github/sokrypton/ColabFold/blob/main/AlphaFold2.ipynb was used for modeling with templates, without amber relaxation and homooligomer set to 1. Comparable models were obtained by modeling with or without templates or amber relaxation. For each SU, the top model with the highest average pLDDT scores was selected for analysis. A 122-amino-acid region in loop G of syncytin-2 SU (aa 158 to 279 from initiation codon) was removed for modeling. Initial modeling of full-length syncytin-2 SU showed a distorted structure with parallel β-strands 5 and 6 significantly displaced. The deleted region in the final model was determined by empirically testing different deletions in modeling to identify the shortest deletion yielding models with the β-strand 5 and 6 homologues in the apical region in the expected location of the PD.

### Model analyses

SU models were analyzed with PyMOL version 2.5.1 (Schrödinger, LLC). The pLDDT scores were extracted from the column corresponding to B-factors (column 11) in the PDB files with atomic coordinates of the models. Secondary structure and disulfides were automatically determined by PyMOL. Structural alignments and similarities between models and with other structures in the Protein Data Bank were determined with the DALI server (ekhidna2.biocenter.helsinki.fi)^42^. DALI similarity Z-scores above 2 were considered significant^42^. The RBD and linker sequences in the SU models of gammaretroviruses, syncytin-1 and syncytin-Mar1 up to 1 residue prior to the CWLC consensus sequence were removed before running structural alignments to avoid alignments of this gammaretrovirus-specific domain.

## Data availability

Atomic coordinates and pLDDT scores in PDB format of the retroviral SU structural models will available at the Dryad database

## Supplementary Tables

**Supplementary Table S1** – Accession numbers, envelope protein sequences and SU sequence boundaries used for modeling.

