## Supplementary Figures S1 to S8 for "Deep time structural evolution of retroviral and filoviral surface envelope proteins"

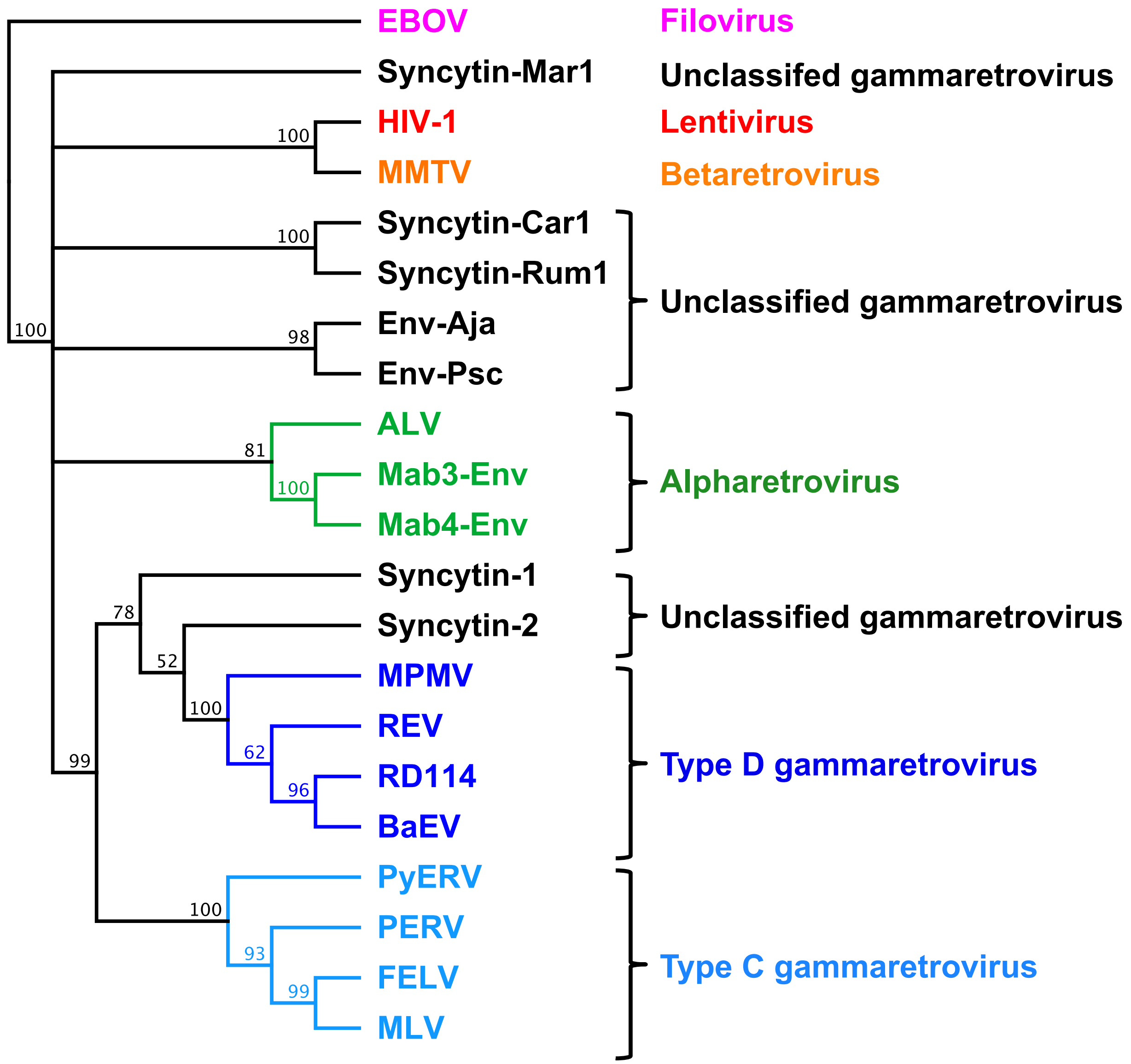


**Supplementary Figure S1** – Relationship of orthoretroviral TM and filoviral $GP_2$ transmembrane subunit ectodomain sequences. A neighbor-joining cladogram was generated with Geneious Prime version 2021.2.2 using Ebola virus (EBOV) $GP_2$ to root the cladogram. Orthoretroviral and filoviral groups are color-coded and indicated next to the cladogram. Numbers indicate node support (percent) from 1000 bootstrap runs.

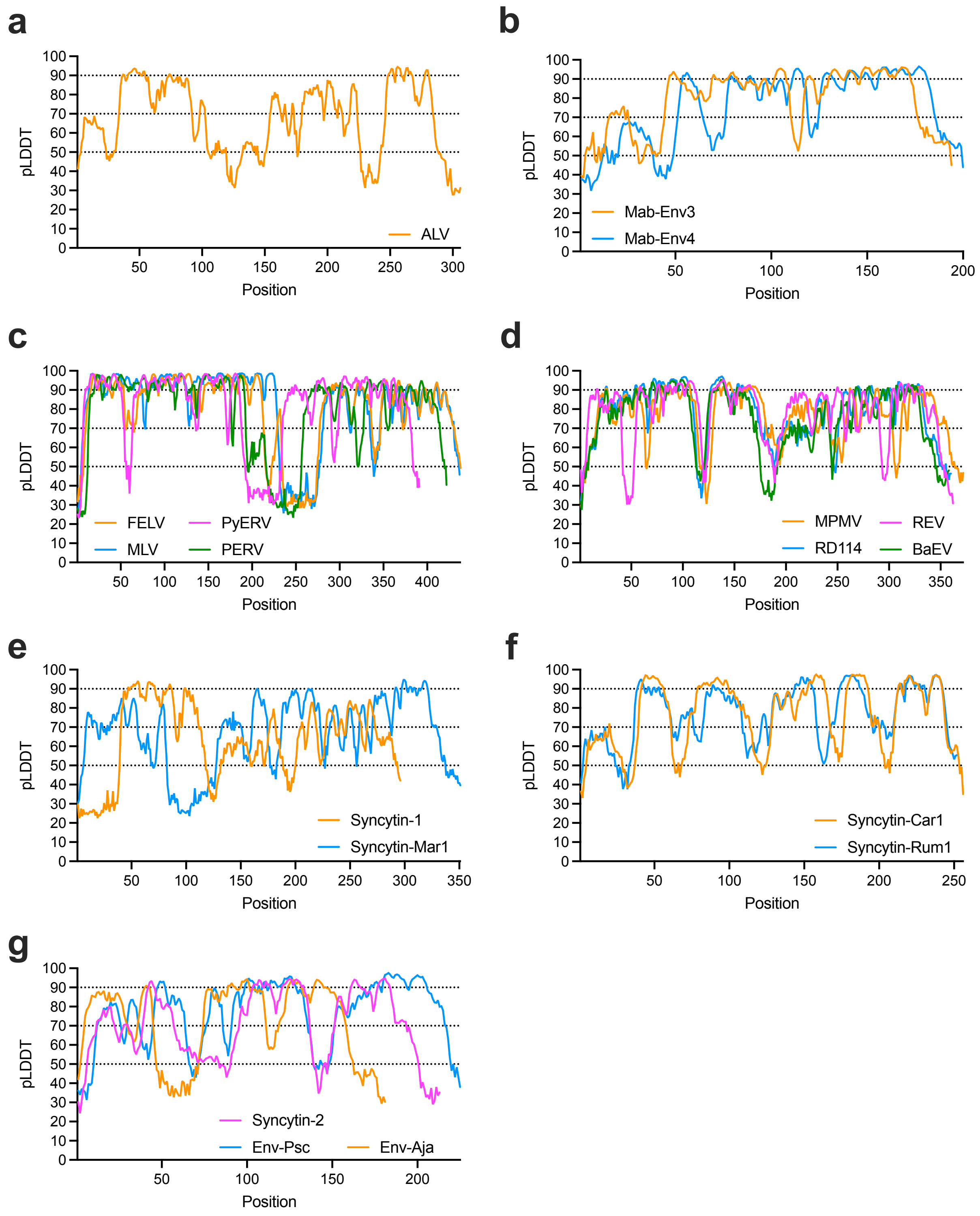


**Supplementary Figure S2 –** pLDDT scores of SU models. pLDDT scores along the SU sequence are indicated for each model, grouping similar models in the same panels.

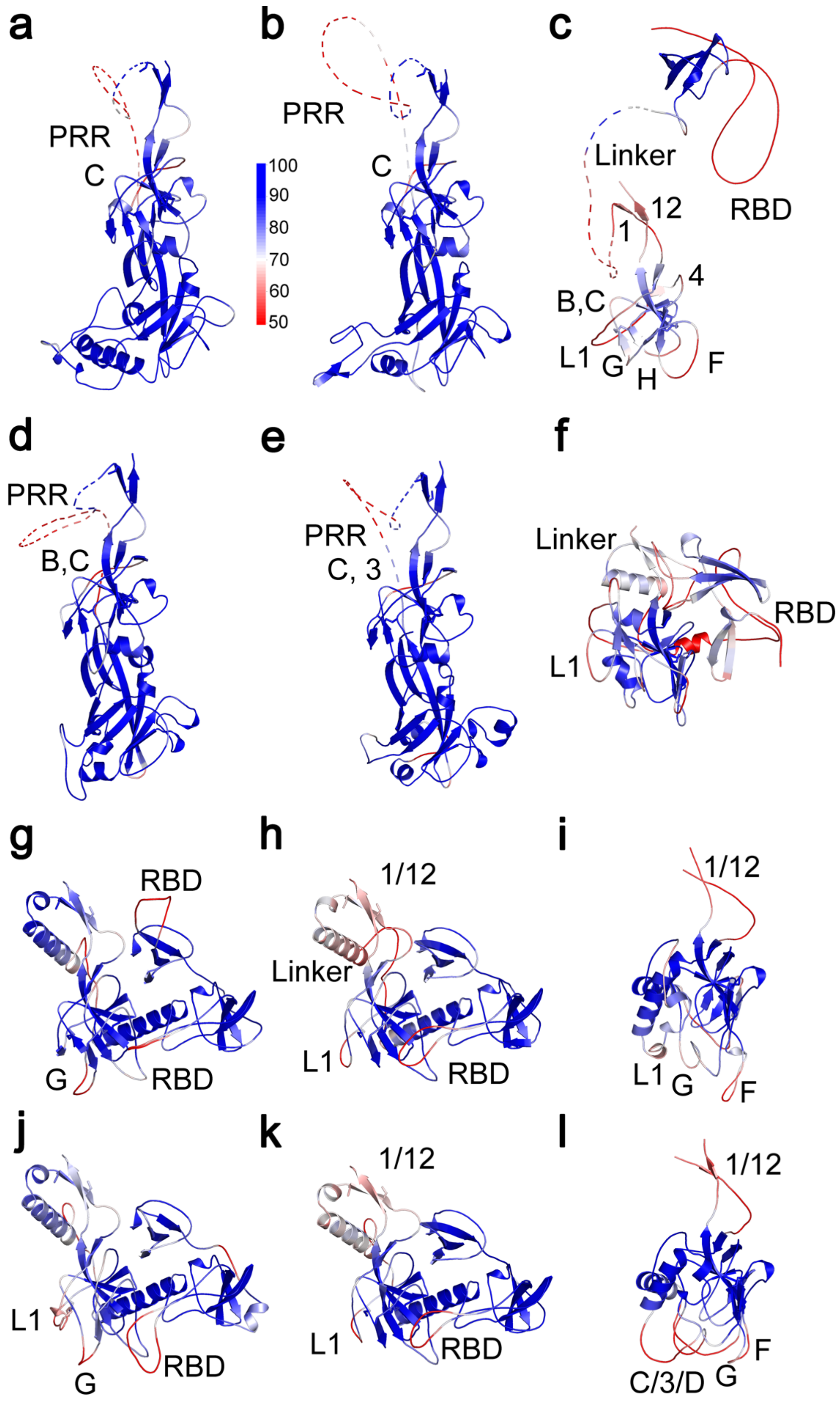


**Supplementary Figure S3 - continued**

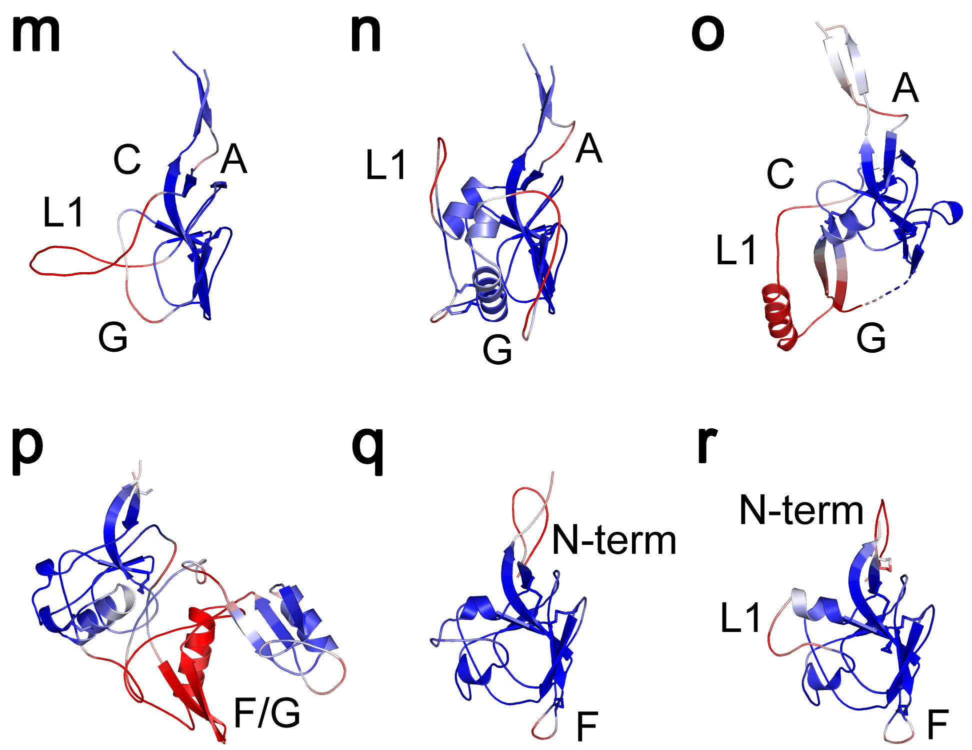


**Supplementary Figure S3** – pLDDT scores of orthoretroviral SU models. pLDDT scores mapped on cartoon representation of (a) MLV, (b) FELV, (c) syncytin-1, (d) PERV, (e) PyERV, (f) syncytin-Mar1, (g) REV, (h) BaEV, (i) syncytin-Rum1, (j) MPMV, (k) RD114, (l) syncytin-Car1, (m) Env-Aja, (n) Env-Psc, (o) syncytin-2, (p) ALV, (q) Mab-Env3, (r) Mab-Env4 SU models. The pLDDT score scale is shown next to panel a. Scores below 50 are shown as red. Unstructured PRR and linker regions are shown as dotted lines. Regions with pLDDT scores below 70 are labeled in each panel.

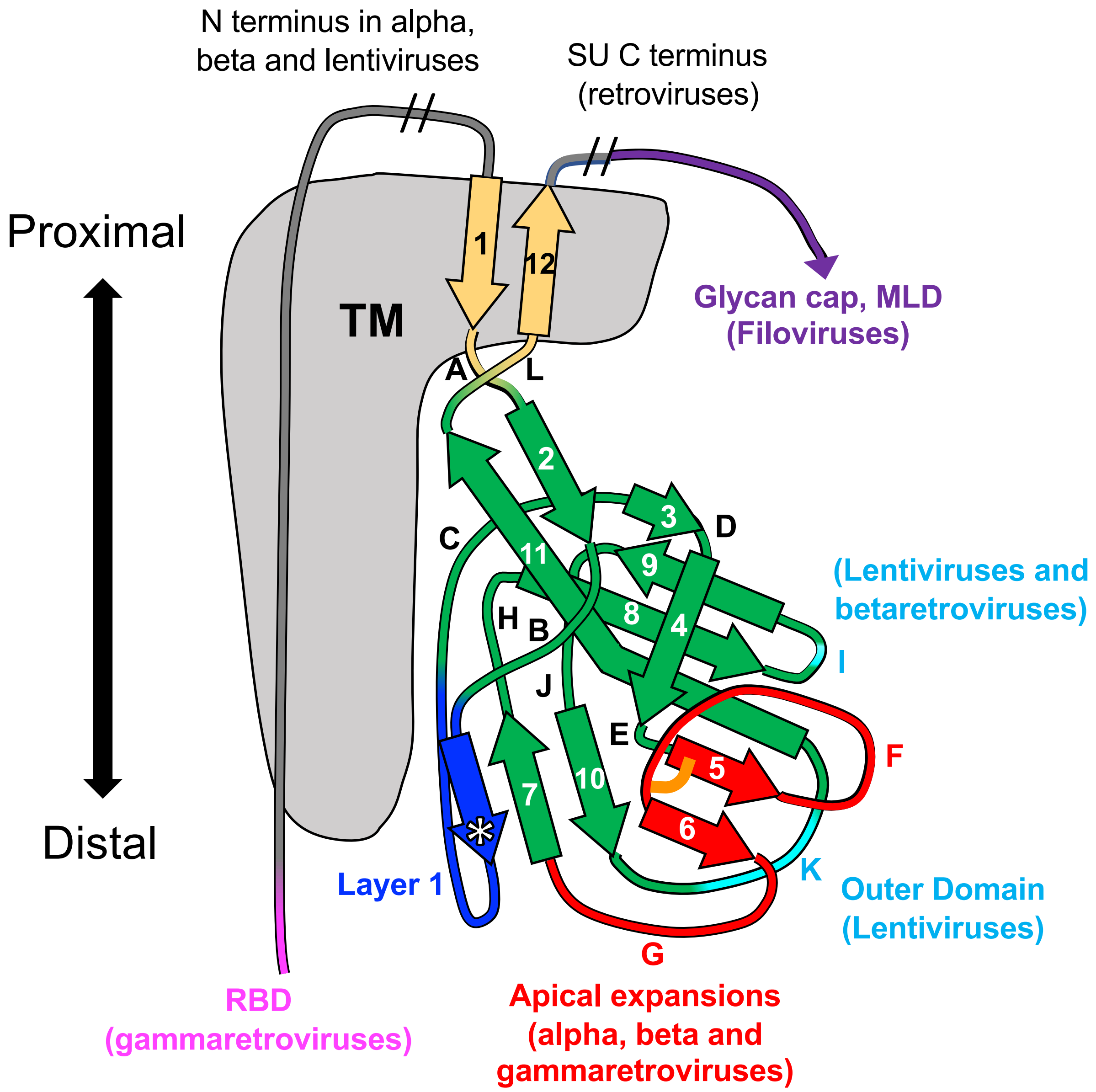


**Supplementary Figure S4 –** Schematic representation of the PD structure and major extended regions in different viral lineages. Regions are colored as in Fig. 3 and 4, with the PD, apical region, layer 1 and K regions shown in green, red, blue and cyan. The approximate location of HIV-1 gp41 in the trimer structure is shown in gray. The asterisk in layer 1 indicates a β-strand often present that extends the β-sheet formed by extended β-strands 7 and 10. The conserved disulfide bond between β-strands 5 and 6 in the apical domain is shown as an orange line.

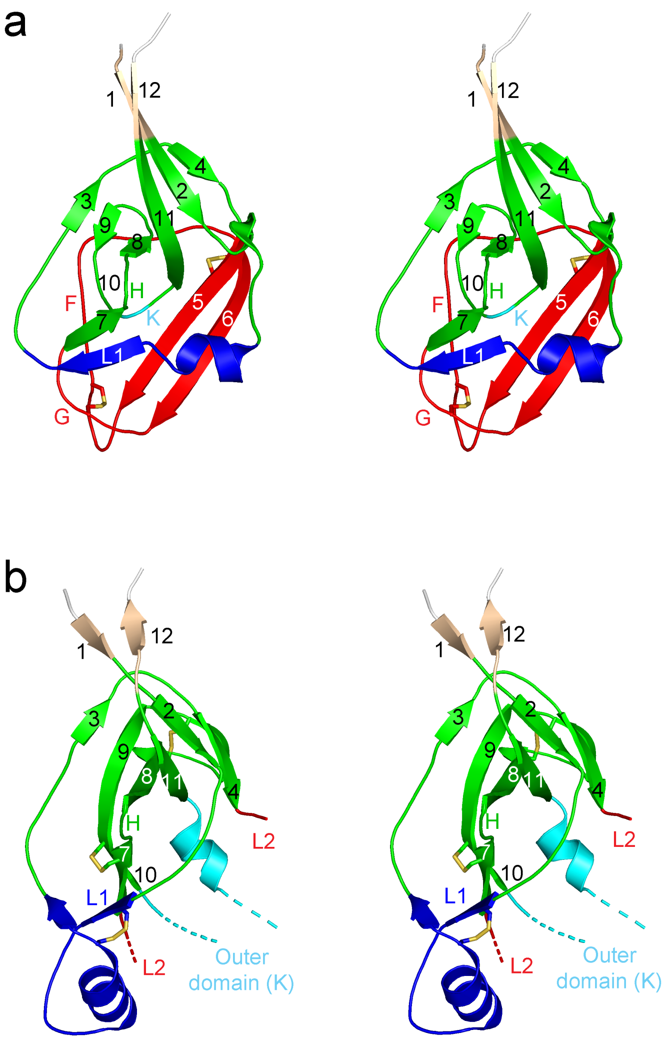


**Supplementary Figure S5 –** Comparison of EBOV $GP_1$ and HIV-1 gp120 PD structures. Stereo view of the PD regions of (a) EBOV $GP_1$ (PDB 3CSY) and (b) HIV-1 gp120 (PDB 3JWD) crystal structures. Both structures are shown in similar orientations with region H region, which in HIV-1 gp120 faces the trimer axis, pointing out of the page. Homologous β-strands and selected loop or expansion regions are shown following the numbering and naming system and color scheme shown in Fig. 3 and 4, with the PD, apical region, layer 1 and K regions shown in green, red, blue and cyan. The extensions forming the distal region, or layer 2 (L2), of the inner domain and outer domain of gp120 topologically equivalent to region K are omitted for clarity. The Glycan cap domain of $GP_1$ following β-strand 12 is not shown. Disulfide bonds are shown as sticks.

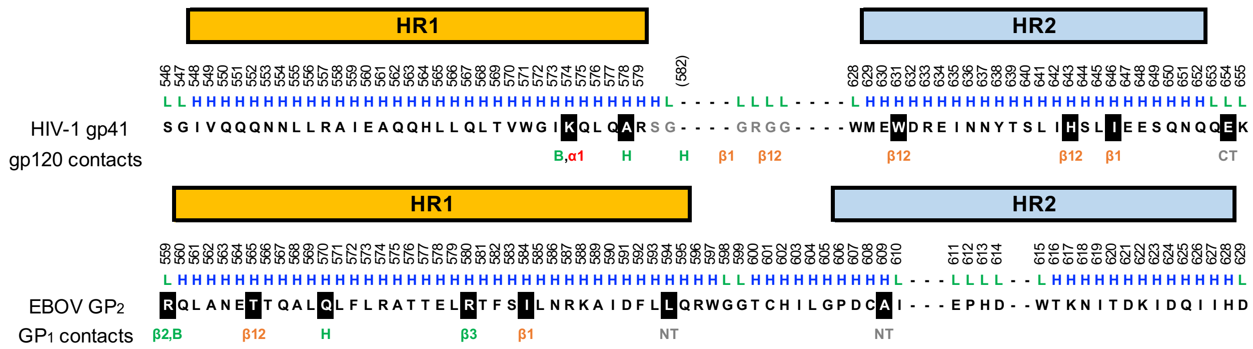


**Supplementary Figure S6 –** Transmembrane gp41 and $GP_2$ subunit contacts with gp120 and $GP_1$ in trimeric envelope protein structures. The sequences of HIV-1 gp41 (PDB 1SZT) and EBOV $GP_2$ (PDB 2EBO) are structurally aligned according to their coiled-coil post-fusion conformation, showing the heptad repeats (HR) 1 and 2. Residue numbering and secondary structure are shown above the sequences. The gp120 and $GP_1$ contact regions for gp41 and $GP_2$ residues in the trimeric, prefusion structures are indicated under the sequences. The region between the two HR in HIV-1 gp41 is replaced with a linker in the structure and the approximate position of contacts in the trimeric structure are shown for that region. Contacts with the fusion peptide regions are not shown.

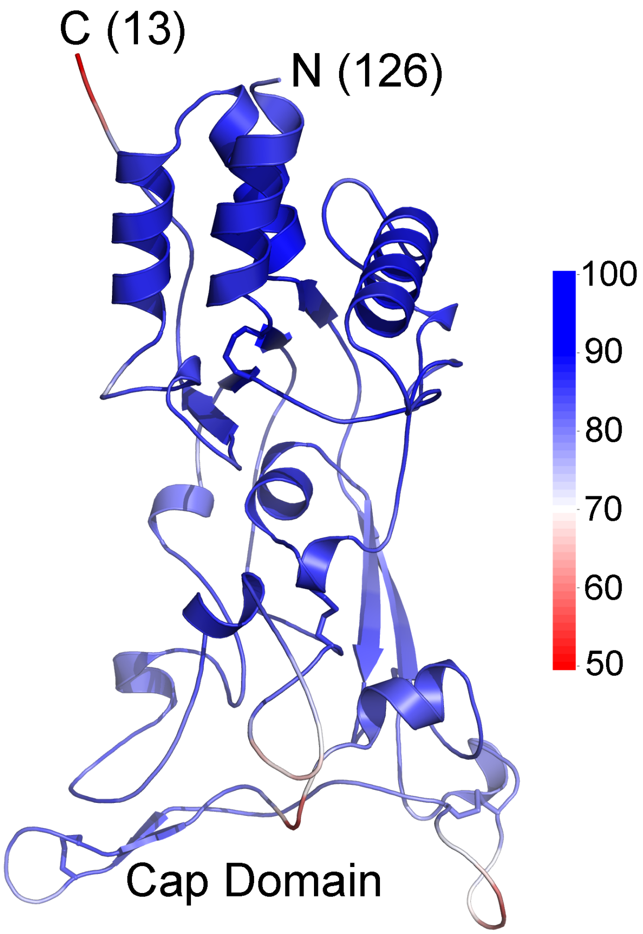


**Supplementary Figure S7 –** African Green Monkey simian foamy spumaretrovirus SU model. The modeled structure is shown with colors indicating pLDDT scores along the chain, with the scale shown to the right. The amino and carboxy termini of the model are indicated with the number of residues not modeled on each end. The amino terminal section outside of the model shown has helical regions that not pack with the rest of SU and are not structurally similar to the SU of orthoretroviruses. Disulfide bonds are indicated by sticks. Models for the SU of other extant spumaretroviruses and reptilian endogenous spumaretroviral elements show the same general structure as the model shown.

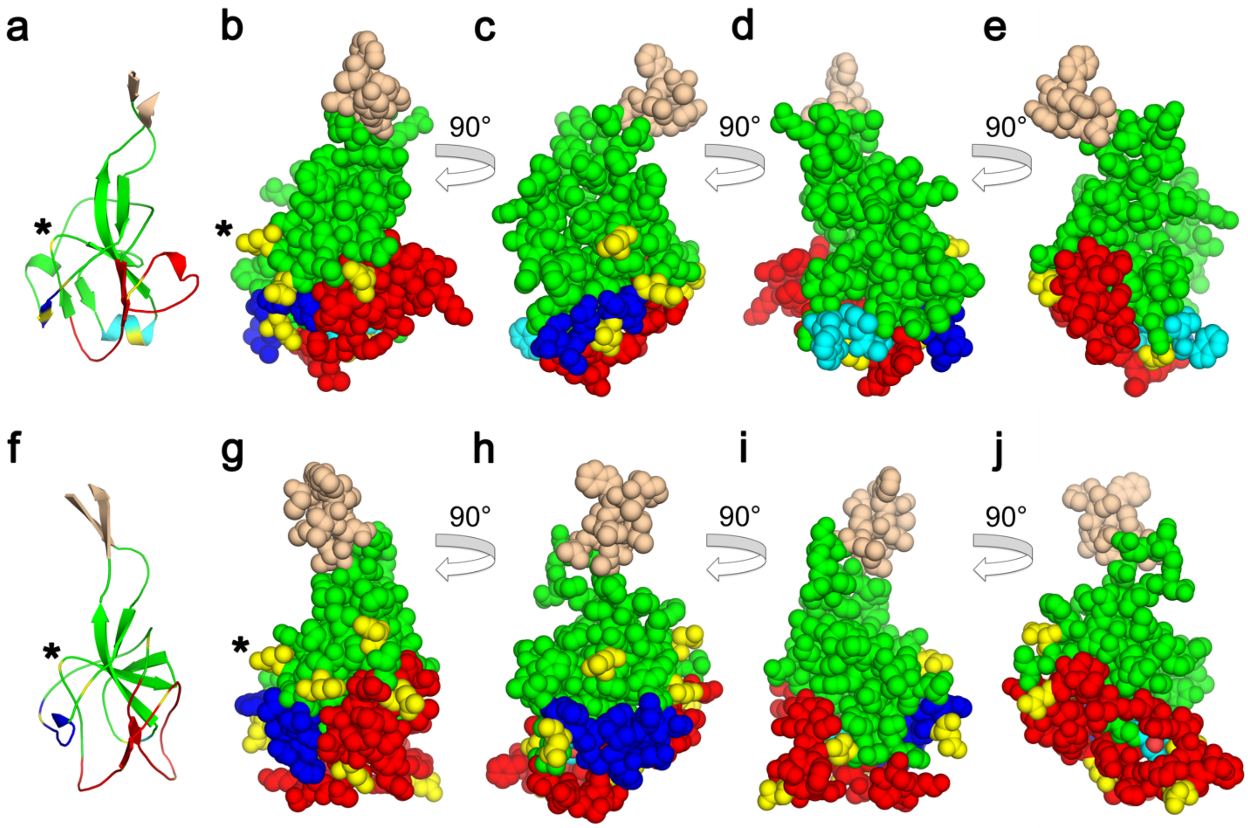


**Supplementary Figure S8** – Glycosylation sites of gammaretroviral PD regions. (a) Cartoon representation of the FELV PD model. (b) Space-filling representation of the same model as in panel a. (c-e) 90° rotations of the model in panel b. (f) Cartoon representation of MPMV PD model. (g) Space-filling representation of the same model as in panel f. (h-j) 90° rotations of the model in panel g. Regions are colored as in Fig. 4, with the PD, apical region, layer 1 and K regions shown in green, red, blue and cyan.. Asparagine residues in potential glycosylation sites are shown in yellow. The H region that in HIV-1 is closely associated with gp41 is indicated with an asterisk. Note the glycosylation site centrally located in region H in panels b, c, g, and h. Regions prior to and beyond $\beta$-strands 1 and 12 are not shown.
